# Turing Test Inspired Method for Analysis of Biases Prevalent in Artificial Intelligence-Based Medical Imaging

**DOI:** 10.1101/2022.05.22.493000

**Authors:** Satvik Tripathi, Alisha Isabelle Augustin, Farouk Dako, Edward Kim

## Abstract

**Background:** Because of the growing need to provide better global healthcare, computer-based and robotic healthcare equipment that depend on artificial intelligence have seen an increase in development. In order to evaluate artificial intelligence (AI) in computer technology, the Turing test was created. For evaluating the future generation of medical diagnostics and medical robots, it remains an essential qualitative instrument.

**Method:** We propose a novel methodology to assess AI-based healthcare technology that provided verifiable diagnostic accuracy and statistical robustness. In order to run our test, we used a State-of-the-art AI model and compared it against radiologist for checking how generalized of the model is and if any biases are prevalent.

**Results:** We achieved results that can evaluate the performance of our chosen model for this study in a clinical setting and we also applied a quantifiable methods for evaluating our modified turing test results using a meta-analytical evaluation framework.

**Conclusion:** This test provides a translational standard for upcoming AI modalities. Our modified Turing Test is a notably strong standard to measure the actual performance of the AI model on a variety of edge cases and normal cases and also helps in detecting if the algorithm is biased towards any one type of case. This method extends the flexibility detect any prevalent biases and also classify the type of bias.

## 1 Introduction

Artificial intelligence is a computer science discipline that can analyze complicated medical data. In many clinical contexts, their ability to exploit a relationship with data collection can be employed in the diagnosis, treatment, and prediction of results [1, 2, 3].

Artificial intelligence systems are computer programs that allow computers to operate in ways that make it appear intelligent. Alan Turing (1950), a British mathematician, was one of the pioneers of modern computer science and artificial intelligence [4]. He characterized that the intelligent behavior in a computer has the capacity to exhibit human-level performance in cognitive activities, subsequently known as the “Turing test” [5, 6]. The Turing Test is one of the most debatable issues in artificial intelligence and cognitive science, as some machines might not pass his test but it may still be intelligent. Alan Turing proposed the Turing Test (TT) in his 1950 Mind article ‘Computing Machinery and Intelligence’ (Turing, 1950) replacing the question “Can machines think?” [7] The goal of Turing’s work is to provide a mechanism for determining whether or not a computer can think. His paper has been seen as the “starting point” of artificial intelligence (AI), whereas the TT has been regarded as its final objective. He further proposes the Imitation Game to give this idea a concrete form [8, 9, 10, 8].

Researchers have been investigating the possible uses of intelligent techniques in every sector of medicine since the last century. Medical AI has witnessed a rise in popularity during the previous two decades. AI systems can consume, analyze, and report vast amounts of data from various modalities to diagnose disease and guide healthcare choices. In addition to diagnosis, AI can aid in the prediction of cancer patient survival rates, such as lung cancer patients. In the field of radiology, artificial intelligence (AI) is being utilized to diagnose disorders in patients using CT scans, MR imaging, and X-rays [4, 11, 12, 1, 13]. Alongside, the question of fairness and ethics has also become very crucial as more and more techniques are getting ready to be implemented in a clinical setting [14, 15, 16, 17].

## 2 Problem Statement

Our prime question in regrads with the advancements of the state-of-the-art AI-based medical imaging algorithms and devices, how do we compute the performance of the algorithim before actually deploying and decide if it is better or at least as good as a clinician in a real-life medical setting or not? Which also raises the concern of can we completely trust an AI and give it the status of an individual entity or do we need a clinician in the loop to oversee the predictions made by the AI algorithm? [18] Also, we can examine if the current state-of-the-art techniques are good enough for clinical use or if we need more advancements in the development by comparing if they have the same level of preciseness and accuracy on a diverse cohor of patients as a professional clinician with years of experience[19].

### 2.1 Aim and Objective

With the rise of AI-based radiological devices and algorithms providing clinical, diagnostic, and prognostic predictions, along with the accuracy we need to look beyond the performance if the model on certain cases and think about whether these modalities are ethically sound and free of biases or not [20]. Therefore, with our proposed test, we can deeply analyze the predictions made by the algorithm and compare them against humans and see if it is safe enough to be implemented in a medical institution while considering the prevalent biases it may have. The project draws its inspiration from A.M. Turing’s classic Turing test. We propose a modified Turing test which serves as a metric to discover the AI-models true performance in the real-life clinical setting and can also help in detecting any possible biases.

## 3 Methodology

### 3.1 Dataset

For this project, we used two different datasets to train and test our dataset. For the training of our models, we used the publicly available Medical Imaging Data Resource Center (MIDRC) - RSNA International COVID-19 Open Radiology Database (RICORD) [21]. In partnership with the Society of Thoracic Radiology (STR) and the American Society of Nuclear Medicine, the MIDRC-RICORD dataset 1a was developed. For all COVID-positive thoracic computed tomography (CT) imaging studies, pixel-level volumetric segmentation with clinical annotations by thoracic radiology subspecialists was performed according to a labeling schema that was coordinated with other international consensus panels and COVID data annotation efforts.

Database 1a of the MIDRC-RICORD is comprised of 120 thoracic computed tomography (CT) images from four international sites, each of which has been annotated with precise segmentation and diagnostic labeling. For our model training process, we employed 120 Chest CT tests (axial series) as input. The data was retrieved using Cancer Imaging Archive [22].

To test our model we used the COVID-19 CT Lung and Infection Segmentation Dataset publically available at Zenodo [23]. This dataset contains 20 COVID-19 CT scans that have been labeled and annotated. The left lung, right lung, and disease areas were labeled by two radiologists and checked by an expert radiologist before being sent to the pathology lab for testing. The dataset completely fits our research interests because of the additional human-annotated segmentation along with the ground truth.

### 3.2 Segmentation Model

Semantic segmentation of the Lung CT scans was performed using a VGG16-Unet model and compared its performance to other models such as UNet, UNet++, UNet3+, and Attention UNet [24, 25, 26, 27], shown in Figure 1. The choice of VGG16-Unet is because of its similarity to U-Net’s contracted layer and its number of parameters is also less than U-Net[28].

**Figure 1:**
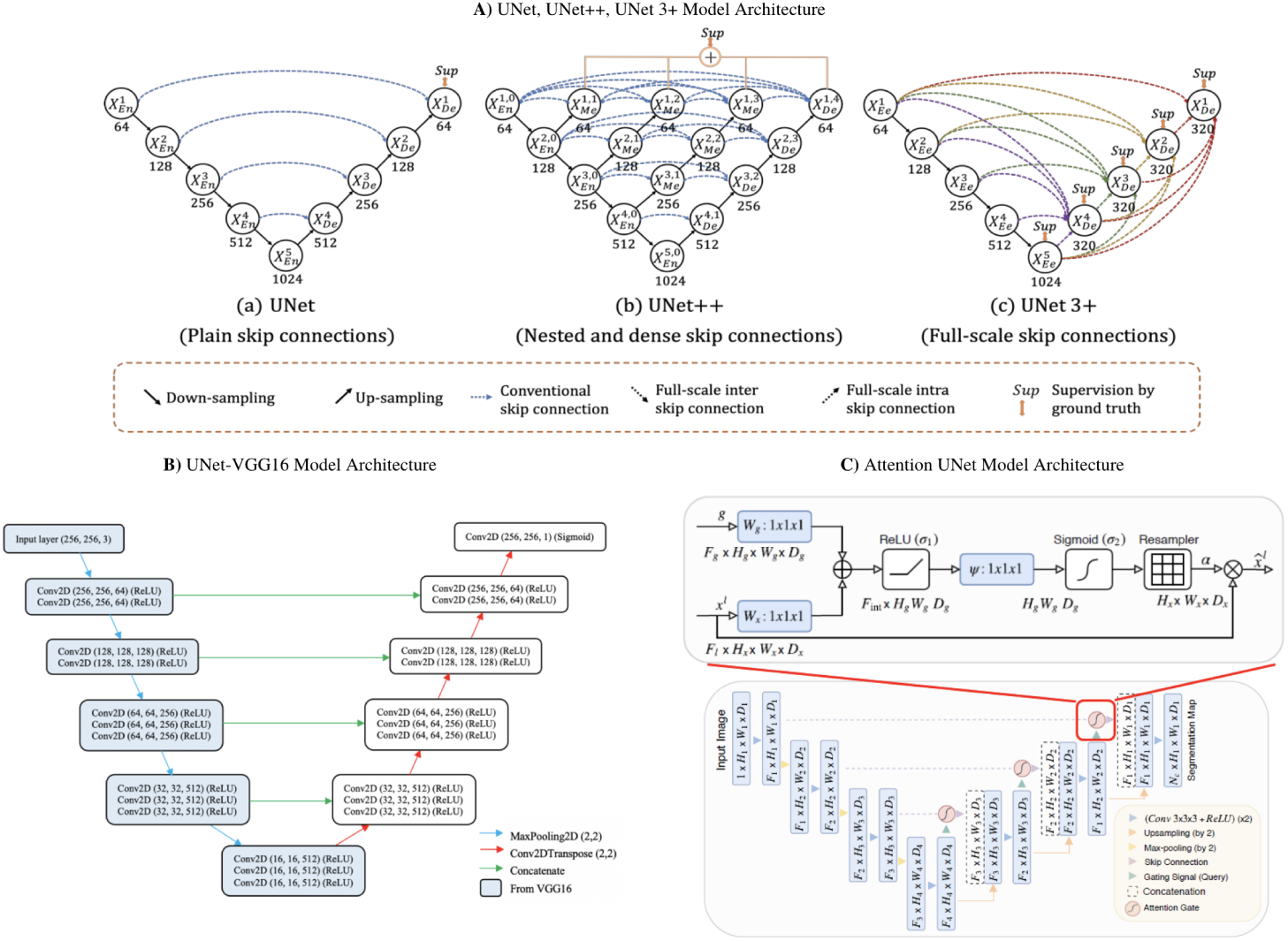
**A)** The model architecture outlining the workflow of UNet, UNet++, and UNet3+. The notable difference between the three models is the skip connections. UNet is using plain skip connections, UNet++ has nested and dense skip connections which have the downside of not being able to explore a sufficient amount of information from full scales. UNet3+, however, uses full-scale skip connections so more information can be obtained during upsampling. **B)** A VGG16-UNet is comprised of an encoder that is based on a VGG-16 model and a decoder that is based on a UNet model. **C)** Attention UNet uses attention mechanisms, compared to a standard UNet model, by focusing on the varying size and shape of target structures.

The left-hand side of the network is an encoder and incorporates the 13 convolutional layers from the original VGG16. After each convolution layer, the MaxPooling operation which reduces the dimensions of the image by 2 × 2 is performed. On the right-hand side of the network, is a decoder. UpSampling operation which restores the dimensions of the image. Each UpSampling operation repeats the rows and columns of the image by 2 × 2. The skip connections are used to restore the dimensions of the image. These skip connections are implemented using the concatenate operation to combine the corresponding feature maps. Since this is a variant of the Fully Convolutional Neural Network, FCN for semantic segmentation, the spatial dimension information of the image needs to be retained hence we use the skip connections. The last convolutional layer has only 1 filter which is similar to a final Dense layer in most other neural networks and gives the binary mask prediction. In total, the network has about 29 convolutional layers which are followed by a PReLU activation. The PReLU has an alpha parameter that is learned during training. In addition, the last convolutional layer has a sigmoid activation function.

The semantic segmentation produced by our proposed UNet-VGG16 is shown in Figure 2. We trained the model on multiple edge cases for producing a more generalized segmentation and the model performed really well in various of these cases.

**Figure 2:**
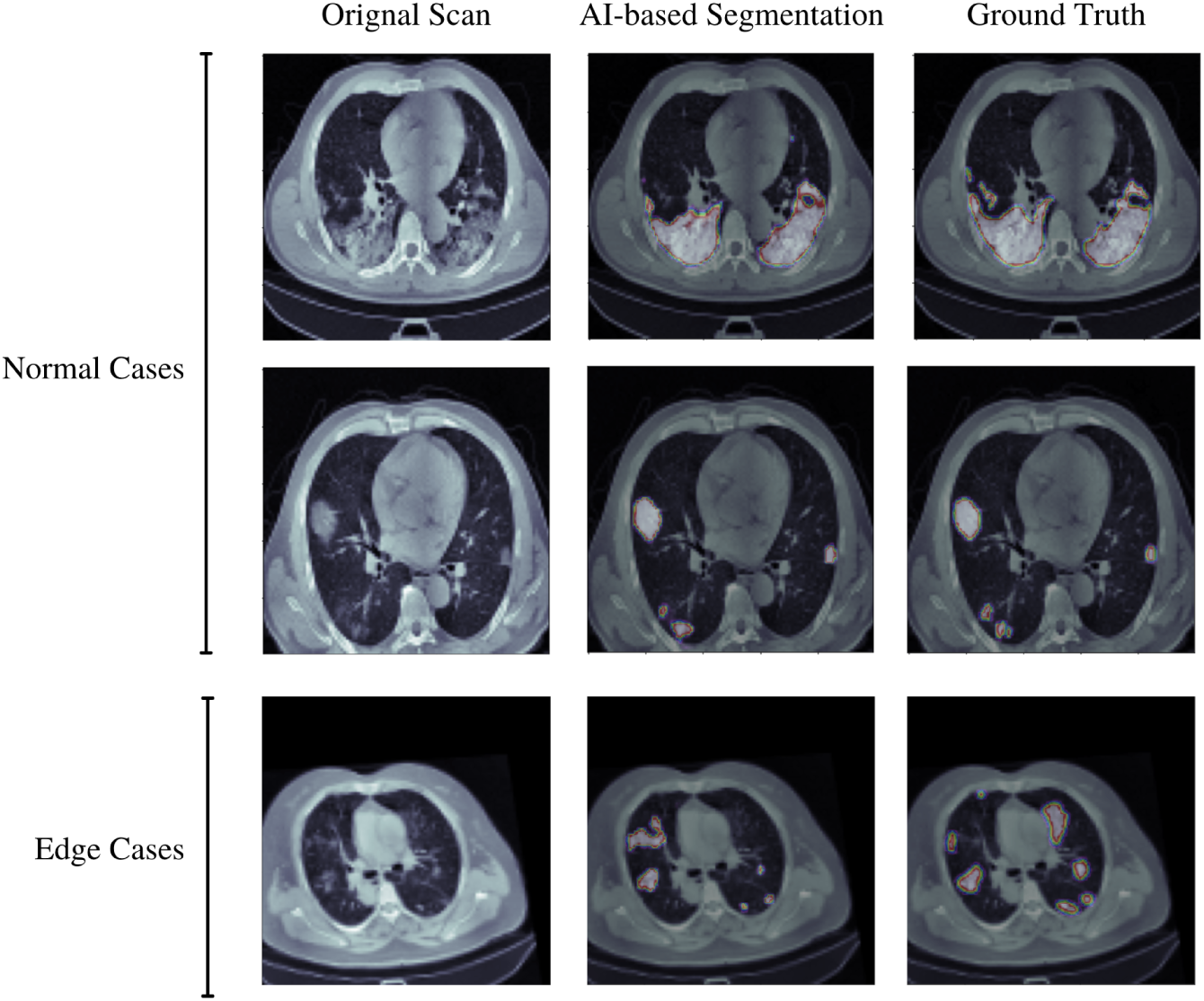
Semantic Segmentation of Normal and Edge Cases Lung Infection Produced by Our Proposed U-Net Model.

### 3.3 Modified Turing Test

This study analyzes the Turing test’s possible usage in healthcare informatics, intending to highlight the broader use of diagnostic accuracy approaches for the Turing test in the present and future AI situations. As a response, we aim to create a model for a measurable diagnostic accurate scoring approach for the Turing test (how distinct are a clinician and AI models?). In diagnostic accuracy testing, we adapted the Turing test to account for false positives and true negatives.

As shown in Figure 4, Examiner (A) (blinded) attempts to differentiate between a human control (B) and a computer test subject (C) versus a human test subject (D). The examiner does not know whether the test subject is human or a machine, therefore (C) vs (D) provides the diagnostic assessment. As a diagnostic test, the redesigned Turing test will now be assessed using a diagnostic accuracy technique and can provide the fast feedback of a human examiner—a method for determining if a computer “(C)” is indistinguishable from its homolog.

The findings of this test may be compared to the results of a gold-standard reference test, namely whether or not the test subject is a computer. The segmentation done by the expert radiologist (D) is shown in figure 3. The radiologist did not see the ground truth while doing the human-annotated segmentation.

**Figure 3:**
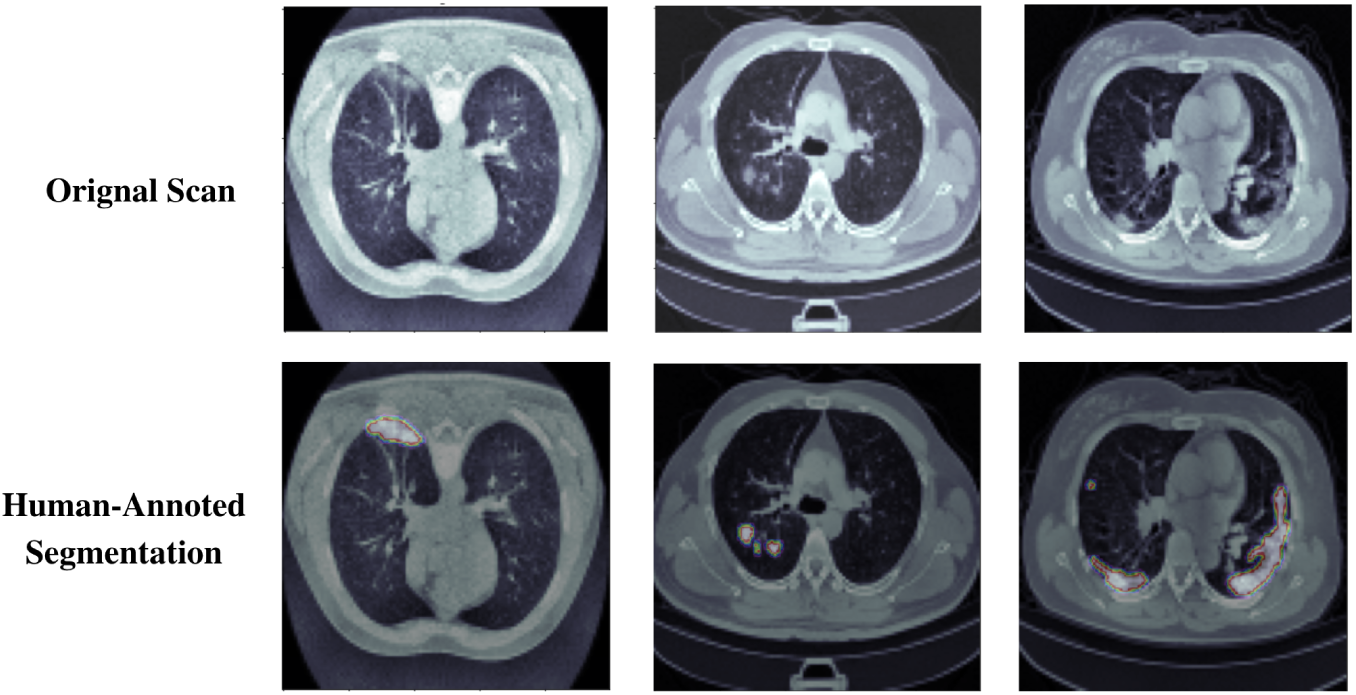
Human-Annotated Segmentation Produced by Expert Radiologist.

**Figure 4:**
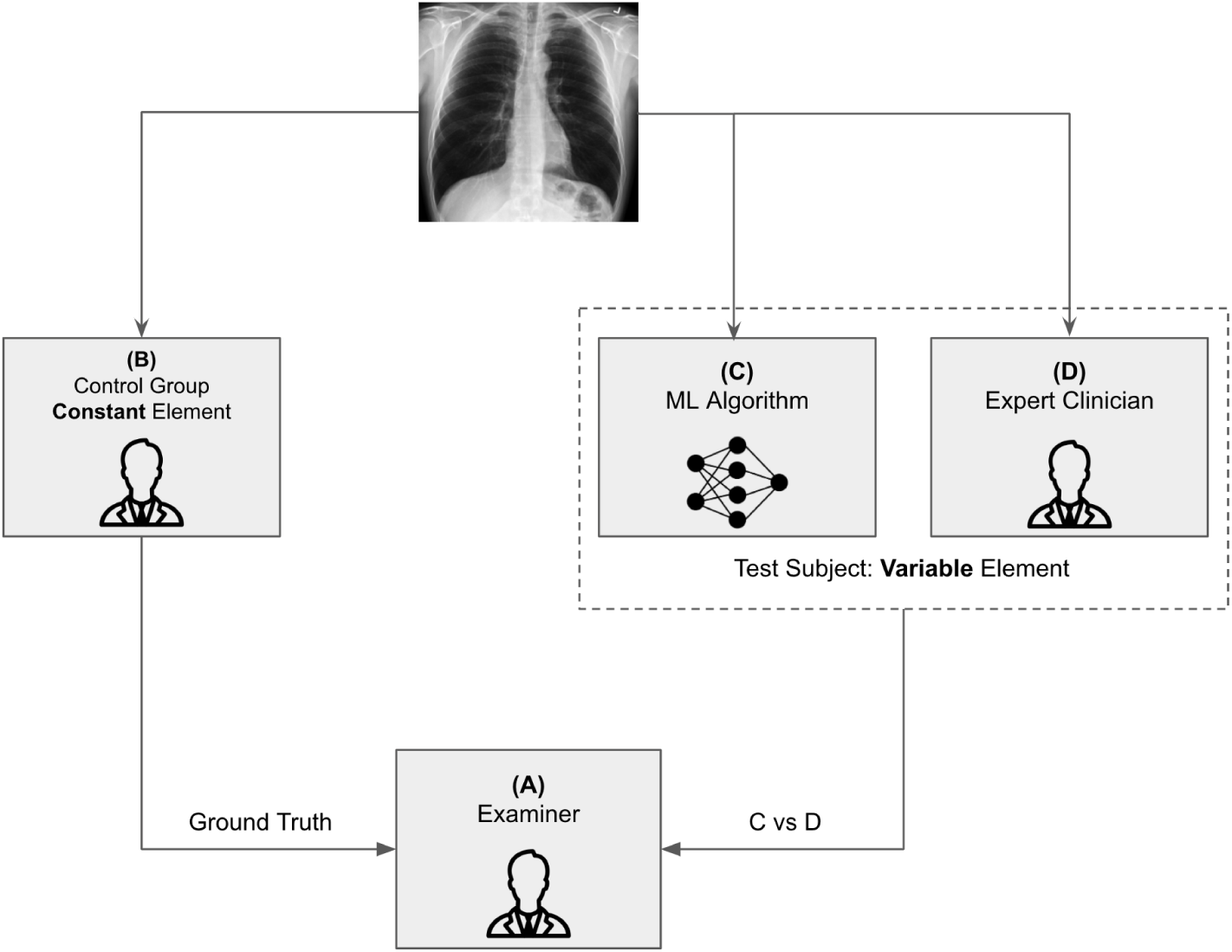
A Systemic Layout of the Modified Turing Test.

The participants (radiologists) of the test were asked to make the prediction on the basis of how accurate the given segmentation is when compared to the ground truth, and based on this the individual may classify whether the segmentation is absolutely accurate and has details and done by a professional radiologist or if it is done by an AI model and it has some missing features. The motive of this study is not to see who does more neat segmentation rather it focuses on whether or not the machine learning algorithms pick up on the clinically important features in the scan.

Consequently, each computer may be evaluated numerous times by the same human and compared to find how biased or accurate the algorithm is. This allows us to obtain several diagnostic evaluations parameters such as sensitivity, specificity, positive predictive value (PPV), and false predictive value (FPV), and we can also generate a receiver operating characteristic (ROC) curve. The proposed diagnostic metrics could be made using the principles of the confusion matrix [29], as shown in Figure 5.

**Figure 5:**
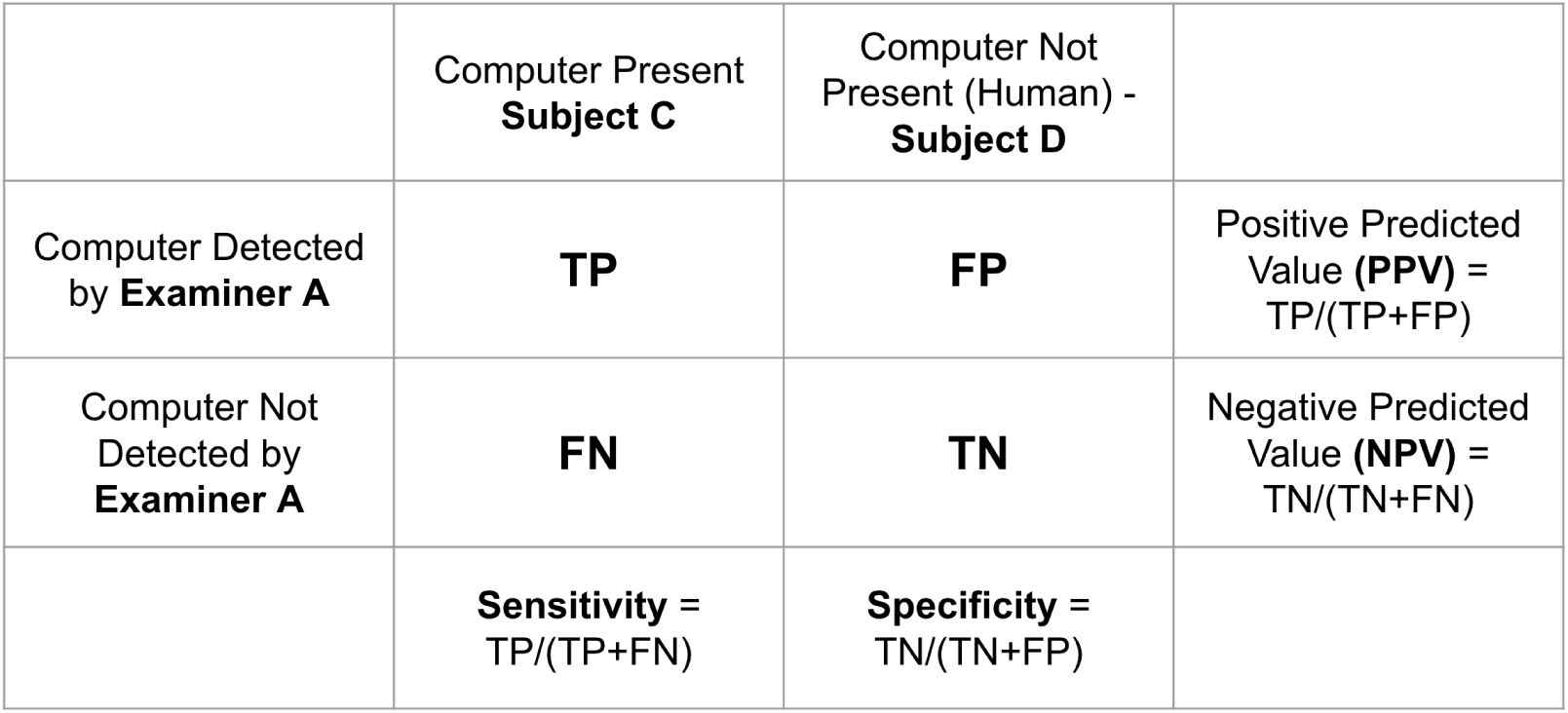
Diagnostic Evaluation Metrics Generated Through Test Results.

The AI model would be considered more accurate and reliable if the AI predictions make the radiologists believe that the segmentation is done by a real human being in terms of preciseness, picking of the area of interest, and if any important considerations are needed in a scenario of an edge case [30, 31].

This is a technique that has never been implemented before and thus is highly novel. The Turing test modification can provide verifiable diagnostic precision and statistical effect–size resilience in the evaluation of AI for computer-based and robotic healthcare and clinical solutions.

## 4 Results and Discussion

### 4.1 Segmentation Results

In this study we used multiple metrics to evaluate the performance of the model: Dice coefficient (DSC), mean Intersection over Union (mIoU), Recall (RE), Precision (PR), Specificity (SP), and F1-score (F1). The expressions of the metrics are described as follows:

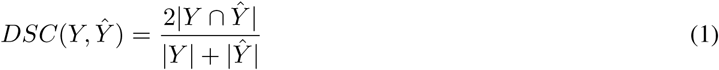

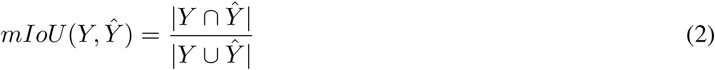

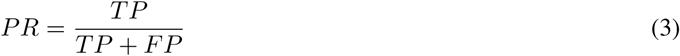

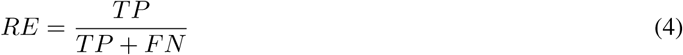

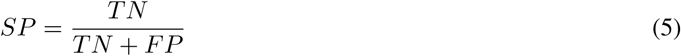

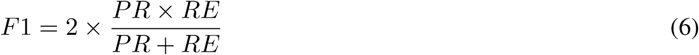

Table 1 compares the segmentation results of the UNet, UNet++, UNet3+, Attention UNet, and UNet-VGG16 models in terms of all metrics used in our experiments. Our proposed Unet-VGG16 model achieved the highest accuracy amongst all other models in all of the matrices.

**Table 1:** Segmentation Metrics Results Amongst Various UNet Models Trained

| Model | DSC | F1 | mIoU | RE | PR | SP |
| --- | --- | --- | --- | --- | --- | --- |
| UNet | 93.42 | 94.65 | 88.36 | 95.31 | 93.28 | 98.71 |
| UNet++ | 93.56 | 94.89 | 87.89 | 94.7 | 93.27 | 97.99 |
| UNet3+ | 95.89 | 96.06 | 88.92 | 95.4 | 94.63 | 98.65 |
| Attention UNet | 94.86 | 95.64 | 88.45 | 95.46 | 94.12 | 98.19 |
| **UNet-VGG16** | **96.73** | **97.94** | **89.21** | **96.98** | **95.56** | **99.41** |

Also, during the testing the model was examined on various edge cases and cases with complex or rare infections to check whether the UNET is biased or not, but the results are very promising, our model achieved a dice score of 94.76% on these critical cases.

### 4.2 Modified Turing Test Results

For this study, 10 board certified radiologists with more than 10 years of experience each in interpreting cardiothoracic imaging reviewed 20 sets of medical images and give out their predictions of whether the segmentation is done by a human or AI based on the preciseness and accuracy of the segmentation. All the radiologist were given the same platform and time to give their predictions. The predictions analysis of each radiologist is given in table 2.

**Table 2:** Analysis of Prediction Derived from the Test Results of the Participants.

| Radiologist | True Positive (TP) | False Positive (FP) | False Negative (FN) | True Negative (TN) | Total Cases |
| --- | --- | --- | --- | --- | --- |
| 1 | 7 | 7 | 4 | 2 | 20 |
| 2 | 1 | 6 | 9 | 4 | 20 |
| 3 | 6 | 7 | 6 | 1 | 20 |
| 4 | 4 | 5 | 8 | 3 | 20 |
| 5 | 2 | 5 | 11 | 2 | 20 |
| 6 | 4 | 7 | 5 | 4 | 20 |
| 7 | 4 | 5 | 10 | 1 | 20 |
| 8 | 5 | 6 | 6 | 3 | 20 |
| 9 | 0 | 7 | 8 | 5 | 20 |
| 10 | 1 | 3 | 12 | 4 | 20 |
| **Total** | **34** | **58** | **79** | **29** | **200** |

The True Positive (TP) denotes that the participant was able to detect the AI-based segmentation and the True Negative (TN) denotes that the radiologist was able to detect the human-based segmentation. False Negative (FN) represents that the participant thought it was an AI while the segmentation was done by a human, whereas in True Negative (TN) the participant thought the segmentation is done by the AI while it was done by a human.

We would consider the TP and TN as our most important metrics here as they reveal the most context about the performance of the AI algorithms in a clinical setting. TP score reveals the reliability of AI-based segmentation, participants reported when the segmented scan did not include not so obvious infections or overdid some of the areas, it made it easier to say for them that the segmentation was done by an AI because a professional radiologist can never do such segmentation [32]. So, having a high TP score is not a good metric for the AI because it means that the segmentation generated are not clinically relevant enough. TN score is what makes an algorithm come closer to an “expert radiologist.” If the model earns more TN scores that means that the AI system is as good as a professional radiologist and is very hard to distinguish whether the segmentation is done by a human or a machine.

To furthermore understand the metrics we have calculated evaluation matrices like Accuracy, Recall, Precision, and others to understand the overall performance and behavior of participants as well as the AI model, the results are shown in Table 3.

**Table 3:** Evaluation Metrics Results Amongst All Participants.

| Diagnostic Evaluation Metrics | Score (%) |
| --- | --- |
| Accuracy | 31.3 |
| True Positive Rate (Recall) | 30.08 |
| True Negative Rate (TNR) | 33.4 |
| False Negative Rate (FNR) | 69.91 |
| False Positive Rate (FPR) | 66.7 |
| Precision | 36.95 |

Overall, our UNet model did exceedingly well in this test, where not only it achieved a high FN score but also received a low TP ratio. This data distributions explains that the model is compatible enough to get implemented in a clinical setting but at the same time there was also a considerable portion of the FP and TN cases, where the participant distinguished between AI and Humans, so taking that into the account we would still need a clinician in the loop to safe-gourd patient care and false prediction making by the algorithm. We also Incorporated 5 out of 20 to be edge cases and the model, and the AI-based segmentation was picked most of the time as an FP.

Finally, we compared the working and performance of the actual Turing Test proposed by Alan Turing and the Modified Turing Test proposed in this study using bivariate meta-analysis [33], and the results are closely similar to what we expected. In Figure 6, we have shown how our modified Turing test works exactly like the original test and even the UNet model’s performance could analyze using the plot.

**Figure 6:**
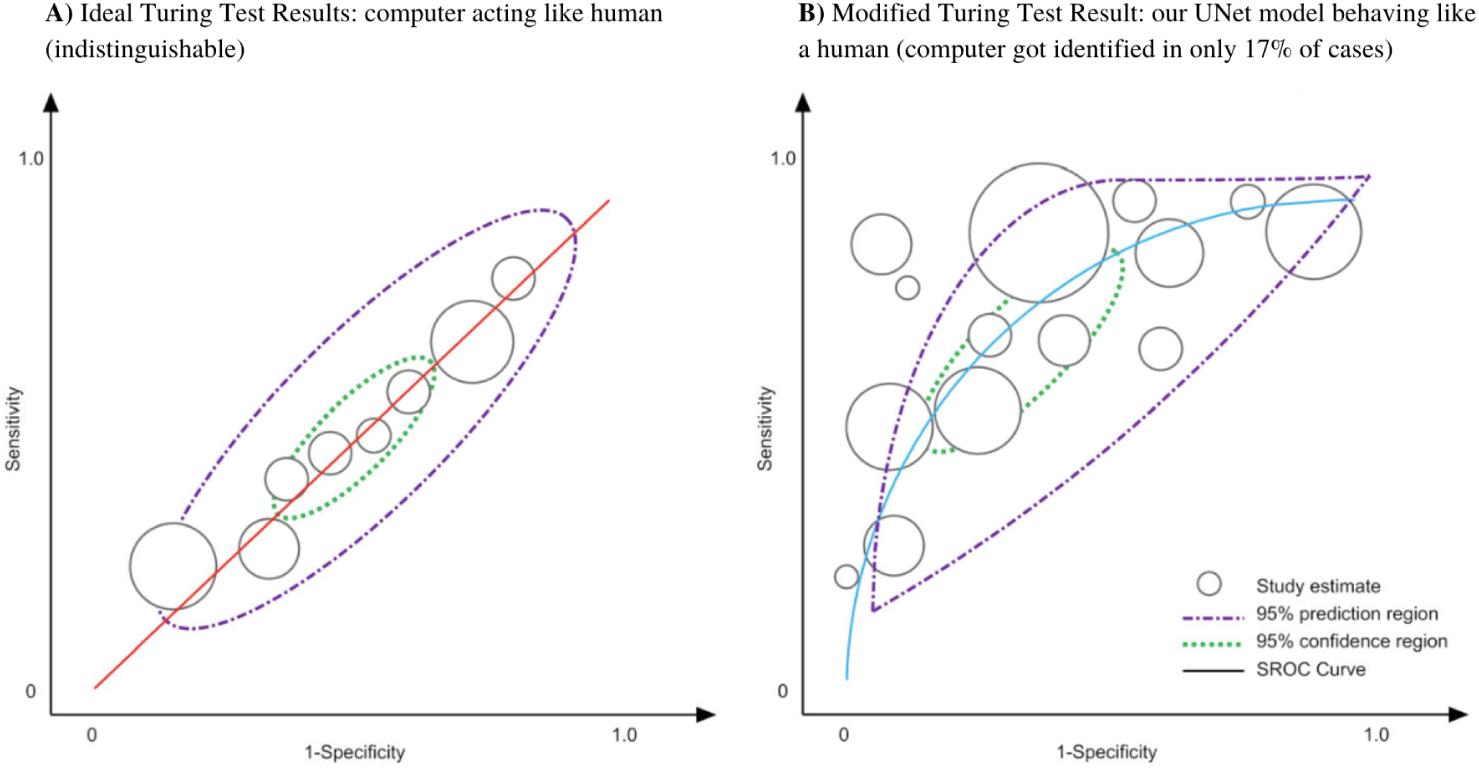
Bivariate Meta-Analysis of Actual Turing Test and Our Modified Turing Test.

## 5 Conclusion and Future Work

The number of AI-based medical imaging devices is getting increased every single day and it is crucial to think about the potential bias it may inherit. This test would be a transnational standard for upcoming AI modalities. We learned through the study we conducted that the use of our modified Turing Test is a notably strong standard to measure the actual performance of the AI model on a variety of edge cases and normal cases and also helps in detecting if the algorithm is biased towards any one type of case. Not just we can detect biases but also classify the type of bias and can work towards resolving it.

Because artificial intelligence systems in healthcare can be utilized for both diagnosis and treatment of diseases, even a tiny error can result in diagnostic inaccuracy and, as a result, increased morbidity and death rates. As a result, it is critical to conduct a comprehensive verification and validation of each artificial intelligence system prior to using it for diagnosis. Consequently, distinguishing between computers and humans (Turing test or modified Turing test) should not detract from the importance of diagnostic accuracy in disease detection and healthcare provision provided by each computer-based AI system, which should be independently appraised for its healthcare safety, precision, and utility. So as we proceed towards the upcoming ages of AI in medicine, this technique would still be applicable in not only segmentation but also in various other prediction and detection models as well. The modified Turing test provides us trust in the AI algorithm and helps us if not look then predict what is inside the black box of the algorithm.

The future of this subject lies in the application of diagnostic accuracy methods to the modified Turing test, which will spur the development of enhanced technology that can closely replicate human behavior in the process of development. This has the potential to produce healthcare computers and other artificial intelligence-based technologies that can improve human health and quality of life while also igniting the next generation of human–technological conversation.

## Acknowledgments

This project is funded by the Drexel Undergraduate Research Mini-Grant. I would like to thank Dr. Brian L Stuart for his guidance and support.

## References

[1] Pavel Hamet and Johanne Tremblay. Artificial intelligence in medicine. Metabolism, 69:S36–S40, 2017.

[2] AN Ramesh, Chandra Kambhampati, John RT Monson, and PJ Drew. Artificial intelligence in medicine. Annals of the Royal College of Surgeons of England, 86(5):334, 2004.

[3] Ahmed Hosny, Chintan Parmar, John Quackenbush, Lawrence H Schwartz, and Hugo JWL Aerts. Artificial intelligence in radiology. Nature Reviews Cancer, 18(8):500–510, 2018.

[4] Satvik Tripathi. Artificial intelligence: A brief review. Analyzing Future Applications of AI, Sensors, and Robotics in Society, pages 1–16, 2021.

[5] Ayse Pinar Saygin, Ilyas Cicekli, and Varol Akman. Turing test: 50 years later. Minds and machines, 10(4):463–518, 2000.

[6] Alan M Turing. Computing machinery and intelligence. parsing the turing test, 2009.

[7] Alan Mathison Turing. Mind. Mind, 59(236):433–460, 1950.

[8] James H Moor. An analysis of the turing test. Philosophical Studies: An International Journal for Philosophy in the Analytic Tradition, 30(4):249–257, 1976.

[9] Gary Marcus, Francesca Rossi, and Manuela Veloso. Beyond the turing test. Ai Magazine, 37(1):3–4, 2016.

[10] Graham Oppy and David Dowe. The turing test. 2003.

[11] Satvik Tripathi, Alisha Augustin, and Edward Kim. Longitudinal Neuroimaging Data Classification for Early Detection of Alzheimer’s Disease using Ensemble Learning Models. 3 2022.

[12] Satvik Tripathi. Early diagnostic prediction of covid-19 using gradient-boosting machine model. arXiv preprint 2110.09436, 2021.

[13] Kun-Hsing Yu, Andrew L Beam, and Isaac S Kohane. Artificial intelligence in healthcare. Nature biomedical engineering, 2(10):719–731, 2018.

[14] Laila Wegner, Yana Houben, Martina Ziefle, and André Calero Valdez. Fairness and the need for regulation of ai in medicine, teaching, and recruiting. In International Conference on Human-Computer Interaction, pages 277–295. Springer, 2021.

[15] Shiri Dori-Hacohen, Roberto Montenegro, Fabricio Murai, Scott A Hale, Keen Sung, Michela Blain, and Jennifer Edwards-Johnson. Fairness via ai: Bias reduction in medical information. arXiv preprint 2109.02202, 2021.

[16] Yoonyoung Park, Gretchen Purcell Jackson, Morgan A Foreman, Daniel Gruen, Jianying Hu, and Amar K Das. Evaluating artificial intelligence in medicine: phases of clinical research. JAMIA open, 3(3):326–331, 2020.

[17] Satvik Tripathi and Thomas Heinrich Musiolik. Fairness and ethics in artificial intelligence-based medical imaging. In Ethical Implications of Reshaping Healthcare With Emerging Technologies, pages 71–85. IGI Global, 2022.

[18] Yoganand Balagurunathan, Ross Mitchell, and Issam El Naqa. Requirements and reliability of ai in the medical context. Physica Medica, 83:72–78, 2021.

[19] Vimla L Patel, Edward H Shortliffe, Mario Stefanelli, Peter Szolovits, Michael R Berthold, Riccardo Bellazzi, and Ameen Abu-Hanna. The coming of age of artificial intelligence in medicine. Artificial intelligence in medicine, 46(1):5–17, 2009.

[20] Onur Asan, Alparslan Emrah Bayrak, Avishek Choudhury, et al. Artificial intelligence and human trust in healthcare: focus on clinicians. Journal of medical Internet research, 22(6):e15154, 2020.

[21] Emily B Tsai, Scott Simpson, Matthew P Lungren, Michelle Hershman, Leonid Roshkovan, Errol Colak, Bradley J Erickson, George Shih, Anouk Stein, Jayashree Kalpathy-Cramer, et al. The rsna international covid-19 open radiology database (ricord). Radiology, 299(1):E204–E213, 2021.

[22] EB Tsai, S Simpson, MP Lungren, M Hershman, L Roshkovan, E Colak, BJ Erickson, G Shih, A Stein, J Kalpathy-Cramer, et al. ‘data from medical imaging data resource center (midrc)-rsna international covid radiology database (ricord) release 1c—chest x-ray, covid+(midrc-ricord-1c). The Cancer Imaging Archive. DOI: https://doi.org/10.7937/91ah-v663, 2021.

[23] Ma Jun, Ge Cheng, Wang Yixin, An Xingle, Gao Jiantao, Yu Ziqi, Zhang Minqing, Liu Xin, Deng Xueyuan, Cao Shucheng, Wei Hao, Mei Sen, Yang Xiaoyu, Nie Ziwei, Li Chen, Tian Lu, Zhu Yuntao, Zhu Qiongjie, Dong Guoqiang, and He Jian. COVID-19 CT Lung and Infection Segmentation Dataset, April 2020.

[24] Olaf Ronneberger, Philipp Fischer, and Thomas Brox. U-net: Convolutional networks for biomedical image segmentation. In International Conference on Medical image computing and computer-assisted intervention, pages 234–241. Springer, 2015.

[25] Zongwei Zhou, Md Mahfuzur Rahman Siddiquee, Nima Tajbakhsh, and Jianming Liang. Unet++: A nested u-net architecture for medical image segmentation. In Deep learning in medical image analysis and multimodal learning for clinical decision support, pages 3–11. Springer, 2018.

[26] Huimin Huang, Lanfen Lin, Ruofeng Tong, Hongjie Hu, Qiaowei Zhang, Yutaro Iwamoto, Xianhua Han, Yen-Wei Chen, and Jian Wu. Unet 3+: A full-scale connected unet for medical image segmentation. In ICASSP 2020-2020 IEEE International Conference on Acoustics, Speech and Signal Processing (ICASSP), pages 1055–1059. IEEE, 2020.

[27] Ozan Oktay, Jo Schlemper, Loic Le Folgoc, Matthew Lee, Mattias Heinrich, Kazunari Misawa, Kensaku Mori, Steven McDonagh, Nils Y Hammerla, Bernhard Kainz, et al. Attention u-net: Learning where to look for the pancreas. arXiv preprint 1804.03999, 2018.

[28] Anindya Apriliyanti Pravitasari, Nur Iriawan, Mawanda Almuhayar, Taufik Azmi, Kartika Fithriasari, Santi Wulan Purnami, Widiana Ferriastuti, et al. Unet-vgg16 with transfer learning for mri-based brain tumor segmentation. Telkomnika, 18(3):1310–1318, 2020.

[29] Olivier Caelen. A bayesian interpretation of the confusion matrix. Annals of Mathematics and Artificial Intelligence, 81(3):429–450, 2017.

[30] Russell T Vought. Re: Guidance for regulation of artificial intelligence applications, 2020.

[31] Daniel L Rubin. Artificial intelligence in imaging: the radiologist’s role. Journal of the American College of Radiology, 16(9):1309–1317, 2019.

[32] Reto Meuli, Yeukuang Hwu, Jung Ho Je, and Giorgio Margaritondo. Synchrotron radiation in radiology: radiology techniques based on synchrotron sources. European radiology, 14(9):1550–1560, 2004.

[33] Hans C Van Houwelingen, Koos H Zwinderman, and Theo Stijnen. A bivariate approach to meta-analysis. Statistics in medicine, 12(24):2273–2284, 1993.

